# A new chapter of the Japanese beetle invasion saga: predicting suitability from long-infested areas to inform surveillance strategies in Europe

**DOI:** 10.1101/2022.11.14.515960

**Authors:** Leyli Borner, Davide Martinetti, Sylvain Poggi

## Abstract

The Japanese beetle (*Popillia japonica*) is a polyphagous pest that spreads rapidly and is estimated to cost more than 460 M$/year in damage and control in the USA alone. This study provides risk maps to inform surveillance strategies in Continental Europe, following the beetle’s introduction and successive spread in the last decade. We developed a species distribution model using a machine-learning algorithm, considering factors relevant to the beetle’s biology, climate, land use and human-related variables. This analysis was performed using presence-only data from native and invaded ranges (Japan, North America, Azores archipelago - Portugal). We gathered more than 30 000 presence data from citizen science platforms and standardized surveys, and generated pseudo-absences using the target-group method. We used the environmental structure of data to randomly sample pseudo-absences, and evaluate model performance *via* a block cross-validation strategy. Our results show that climate, in particular seasonal trends, and human-related variables, are major drivers of the Japanese beetle distribution at the global scale. Risk maps show that Central Europe can be considered as suitable, whereas Southern and Northern European countries are at lower risk. The region currently occupied is among the most suitable according to our predictions, and represents less than 1% of the highest suitable area in Europe. A major cluster of high suitability areas is located near the currently infested zone, whereas others are scattered across the continent. This highlights the importance of designing surveillance strategies considering both active insect dispersal and the possibility of hitchhiking to reach distant areas.

## 1 Introduction

Biological invasions, i.e. successful introduction and establishment of species outside of their native range, are a major issue because of their potential impact on ecosystems (Bellard et al., 2016; Kumschick et al., 2015), food security (Oerke, 2006; Ogden et al., 2019) and economic balances (Diagne et al., 2021). Biological invasions have been greatly facilitated by globalization because movements of goods and people offer countless opportunities for the spread of pests across different regions of the world (Hulme, 2009). The Japanese beetle (*Popillia japonica* Newman; Pj hereafter) is a prime example of a species that successfully reached and established outside its native range through human-mediated transports, resulting in high costs for damaged crops, surveillance and control measures. This species is native to Japan (Clausen et al., 1927) and was first detected outside of its native range when individuals, most likely from Gunma, Ibaraki and Tochigi Prefectures on the Honshu island (Strangi et al., 2022), were found in the United States of America in 1916 (Fleming, 1972). Since this accidental introduction, the beetle has spread throughout North America and Canada. It was then introduced to the Azores archipelago on Terceira island in the 1970’s (Martins et al., 1988; Vieira, 2008) and has spread to all of the Azores islands with the exception of Santa Maria. More recently, Pj has been detected in Northern Italy in 2014 (EPPO Global Database, 2014) and Switzerland in 2017 (EPPO Global Database, 2017). Other interceptions have been observed in Central Europe, while stable populations are only found in the Italian regions of Lombardy and Piedmont and the Ticino region in Switzerland (Poggi et al., 2022).

This highly polyphagous beetle feeds on more than 400 host plants throughout its whole life cycle (EFSA, 2019; Tayeh et al., 2022) and damage costs are estimated at 460M$/year in the USA alone (USDA-APHIS, 2015), while recent studies estimates expected overheads for major EU crops between 30M€/year and 7.8B€/year (Straubinger et al., 2022). European plant health agencies classified Pj as a high priority pest (EFSA, 2019), and Sanchez et al. (2019) found Pj to be the second most important quarantine organism, in terms of economic, social and environmental impact. The challenge is to counteract this invasion at a very early time point to significantly enhance the chances of a successful eradication in already-invaded areas, and containment in uninvaded ones. This brings the design of surveillance strategies to the forefront, the effectiveness of which is crucial as management becomes disproportionately more costly and difficult with increasing incidence of pathogens (Parnell et al., 2014). For this purpose, different authors advocate for the use of risk-based strategies based on the examination of risk maps, generally preferable to systematic sampling programs leading to the suboptimal deployment of resources allocated to surveillance (Hyatt-Twynam et al., 2017).

The objective of the present work is to draw a reliable map of environmental suitability for the Japanese beetle and to discuss implications in view of the design of surveillance and containment strategies, with a special focus on Continental Europe, where a recent invasion is ongoing. Pj suitability is estimated using a species distribution model (SDM) which allows to explore the contribution of a large set of variables describing Pj biology, climate, land use and anthropogenic factors, extracted at a fine spatial resolution (4km x 4km). The SDM is trained using a classification machine-learning algorithm based on presence-only data from native and long-invaded ranges (Japan, North America, the Azores archipelago) and the predicted suitability is projected to the entire Northern Hemisphere. More than 30 000 presence data have been gathered from standardized surveys and citizen-science platforms, whereas the target-group method (Phillips et al., 2009) has been used to generate pseudo-absences in the sake of balancing the presence-only dataset and to account for the sampling bias inherently associated to opportunistic citizen-science data. Finally, we structured the dataset into environmental blocks and sampled pseudo-absences within blocks using environmental distance to presence sites to enforce representation of diverse environmental conditions and to ensure independence of validation data in the cross-validation procedure.

The agreement of our environmental suitability maps with current knowledge of the invasion in long-invaded regions and official infestation status (NPB, 2016) is reported in detail. We discuss how our projections in Continental Europe can contribute to European policymaking and strategies regarding the sustainable management of the invasive Japanese beetle. We conclude with a call for urgent and proportionate surveillance at continental level.

## 2 Materials & Methods

This section describes all data collected and methods used to fit a species distribution model for the Japanese beetle and subsequently predict a suitability index over the Northern Hemisphere and particularly Europe. All data processing and analyses were carried out using R version 4.1.2 (R Core Team, 2021).

### 2.1 Japanese beetle presence data

We collected geo-referenced Pj occurrences from both citizen-science platforms and standardized surveys conducted by phytosanitary services (Figure 1A and Supplementary Information 1). For opportunistic citizen-science observations, data sources and pre-processing are described in the Supplementary Information 1. We also gathered ∼10 000 presence data from standardized surveys conducted by local authorities and phytosanitary agencies in the Azores, Canada, Italy and Switzerland. We did not consider absence data reported by these surveys since the Pj invasion is ongoing in these areas and absence data are unlikely to reflect real absences (Cianfrani et al., 2010). Even if the first record of Pj occurrence dates back to August 1917 in New Jersey (Supplementary Information 1, Figure 1), we considered only presence data starting from 2010 to match the timeframe of most of environmental predictors, hence removing 8% of the presence data before that period.

**Figure 1 :**
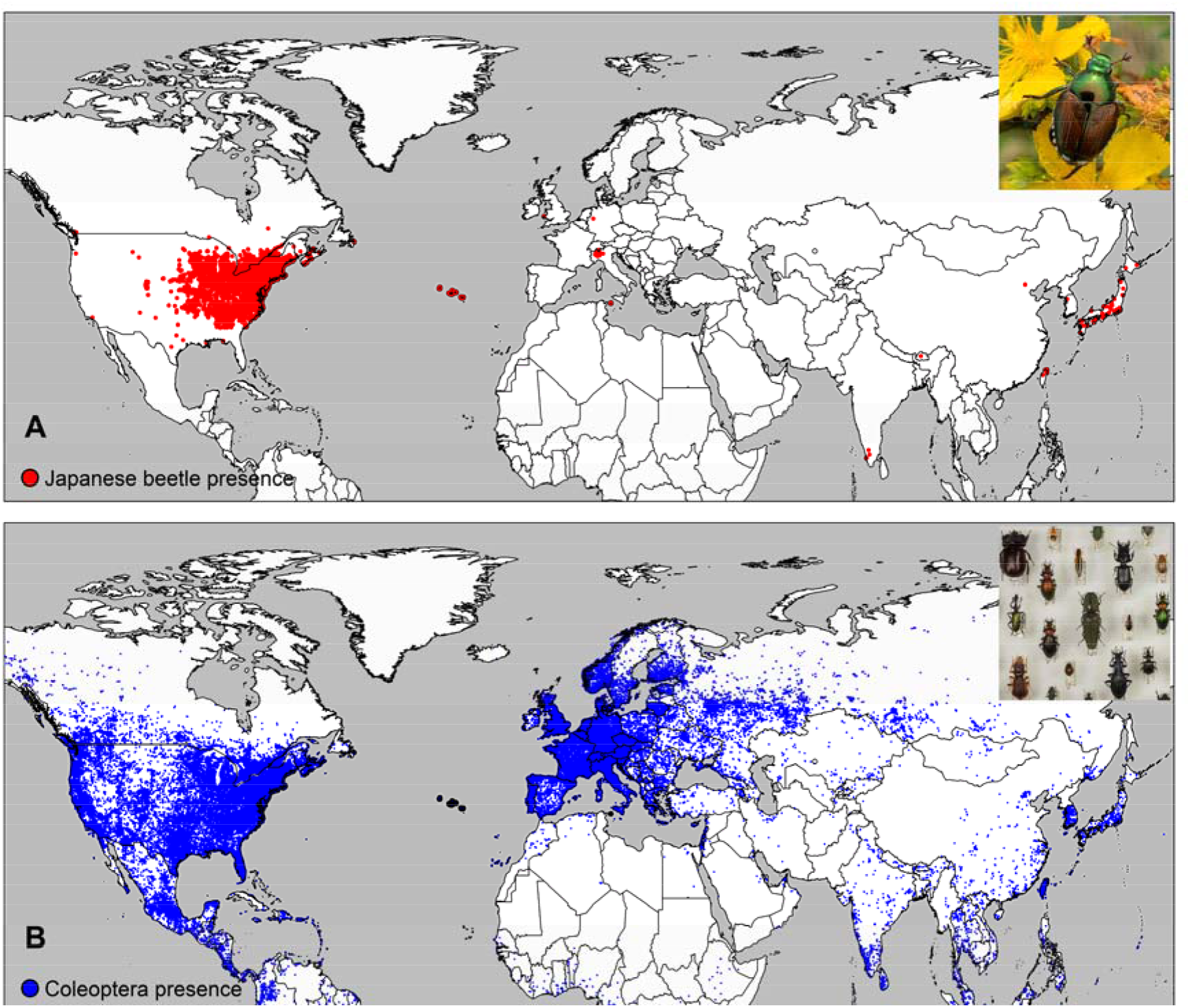
A) Worldwide distribution of geo-referenced Japanese beetle presence data (in red) collected from citizen science platforms and standardized surveys over the period 1917-2021; B) Worldwide distribution of Coleoptera species (target-group, in blue) extracted from the GBIF citizen science platform over the period 2010-2021. Insert photos: A) adult individual of Popillia japonica, © Leyli Borner, INRAE, B) Coleoptera species from Franck Duval’s private collection, © Stéphane Jumel, INRAE

### 2.2 Pseudo-absence data

When confirmed absence observations are not available, one option is to generate pseudo-absences from those locations where presences have never been recorded, but this operation has to be carefully tuned since false absences can negatively affect the SDM (Gu and Swihart, 2004). In general, it is recommended to sample pseudo-absence data within the same range of environmental conditions as in the presence data (Barbet-Massin et al, 2012). Furthermore, due to the origin of our presence data (mainly opportunistic citizen-science observations), pseudo-absences should also be sampled from a set of data that suffer from the same sampling bias as presence data: this technique, known as the target-group method, tends to improve SDM predictions (Phillips et al., 2009). Here we considered all observations of *Coleoptera* (excluding *Popillia japonica*) collected from 2010 to 2021 from the GBIF citizen science platform, for a total of 714 816 observations (Occdownload Gbif.Org, 2021) (Figure 1B).

### 2.3 Predictor variables

More than 300 variables were collected from different data sources to describe environmental conditions associated with the presence of Pj (Supplementary Information 2). These predictors encompass climatic variables, soil characteristics, land uses, and human-related features such as population and road densities. We considered these variables because of their relevance to the Japanese beetle’s biology (Fleming, 1972; Kistner-Thomas, 2019; Zhu et al., 2017), and we did not restrain our analysis to a limited set of variables identified as *a priori* important. Source data had different spatial resolutions ranging from 20 m to 4 km (Supplementary Information 2). We aggregated monthly climatic variables at the seasonal scale (Spring: March to May, Summer: June to August, Fall: September to November, Winter: December to February), and removed variables for which a very high pairwise correlation was found. Eventually, a reduced set of 133 variables was finally retained.

### 2.4 Spatial data aggregation & data availability

In order to harmonize the dataset, we upscaled all data to a 4 km x 4 km resolution (the coarser resolution amongst the environmental predictors), averaging values contained within each cell when necessary (Bierkens et al., 2000). The aggregation rule for presence and pseudo-absence data was defined as follows: a grid cell is labelled as “presence” if at least one observation of the Japanese beetle data is found within the cell, whereas it is considered as “absence” if at least one *Coleoptera* and no Japanese beetle have been reported in the cell. This results in 6844 presence cells and 49010 pseudo-absence cells. The entire dataset, including presence/pseudo-absence data and environmental variables, is available in raster format at a 4km x 4km resolution (Borner et al., 2022).

### 2.5 Modelling species distribution using a machine learning approach

Species distribution model rely on the hypothesis that the modelled species is in equilibrium with its environment (Hutchinson, 1957). Therefore, the species distribution used in model fitting must reflect its realized ecological niche, i.e. the environmental conditions in which a species persists in nature (Guisan & Thuiller, 2005). In the case of biological invasions, it is recommended to use data from both the native and invaded ranges, as shifts in species niche can occur during the invasion process (Barbet-Massin et al., 2018; Broennimann & Guisan, 2008; Guisan et al., 2014). However, in recently invaded ranges, species distribution is likely to be impacted by dispersal limitations and pseudo-absence data sampled in these areas may reflect not-yet-reached areas and not necessarily truly unsuitable conditions (Elith et al., 2010; Liu et al., 2020; Thuiller et al., 2005). Thus, we intentionally excluded data from Continental Europe during model fitting because they represent a recent and ongoing invasion process (less than 10 years since first detection) and considered only observations from either a native or long-occupied invaded area, i.e. Japan, North America (USA and Canada) and the Azores Archipelago. Furthermore, for cross-validation purposes, we trained the SDM only on data from the 2010-2019 period, leaving the remaining data in a separate dataset for validation. In total, 53210 raster cells (4 km x 4 km) are available for model fitting, of which 4200 represent presence data (c.a. 7%), and 49010 represent pseudo-absences (Figure 2 panel A-B).

**Figure 2 :**
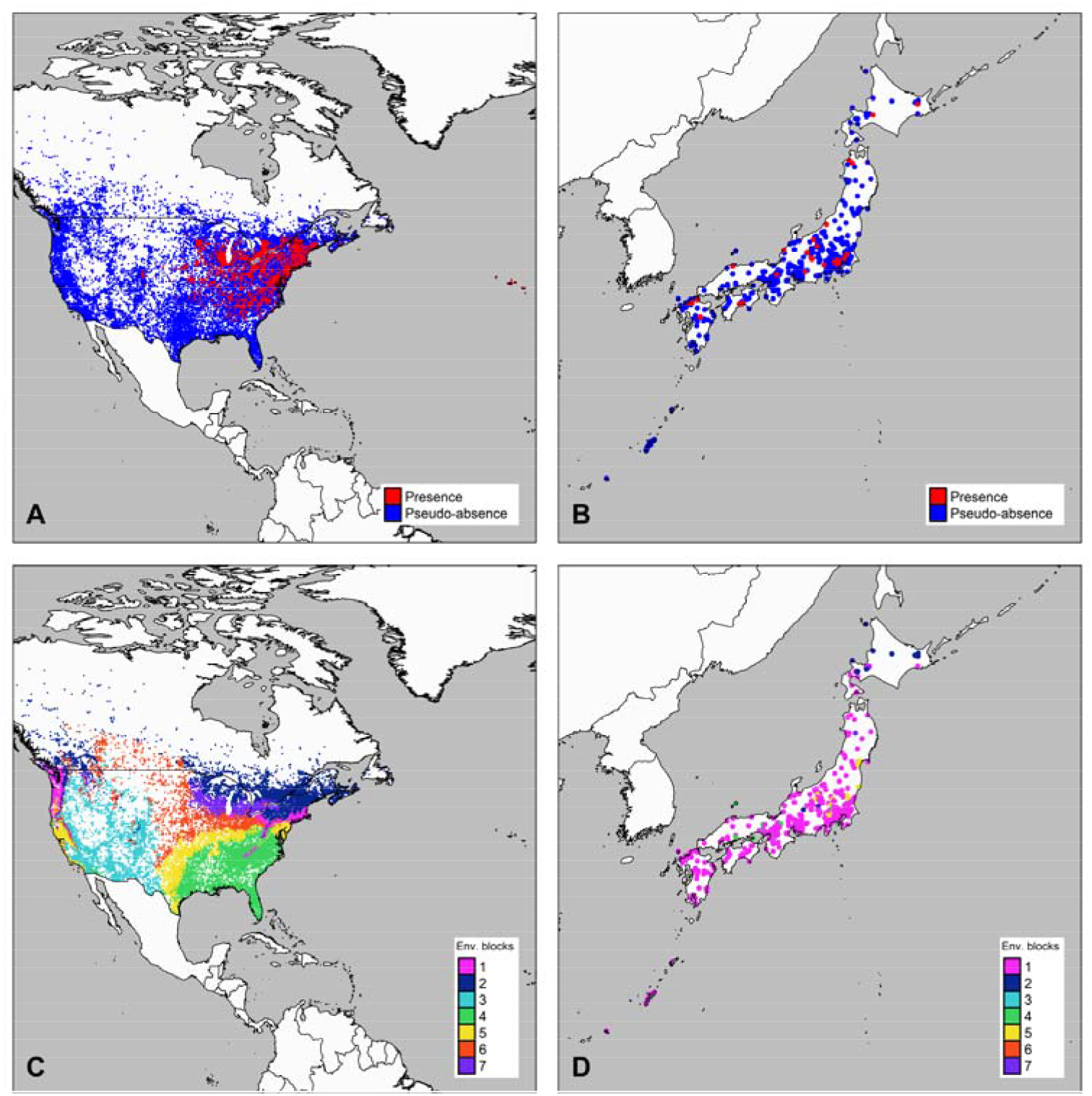
Distribution of Japanese beetle presence data (in red) and pseudo-absences (Coleoptera target-group, in blue) in North America (Panel A) and Japan (Panel B), as well as according to the environmental blocks (N=7) in North America (Panel C) and Japan (Panel D).

Machine learning provides an efficient framework to develop predictive models such as SDM. In this study we fitted a random forest model (hereafter RF) as it has been shown to be a good candidate for species distribution modelling and to be more robust against collinearity among predictors (Cutler et al., 2007; Valavi et al., 2022; Zhang et al., 2019). RF is a recursive partitioning method that is widely used for nonlinear classification. It involves an ensemble of classification trees fitted on bootstrap samples of the data, using a random restricted subset of the predictors for each split in each tree to avoid correlation between trees. The RF algorithm also provides a measure of variable importance (Breiman, 2001) by measuring the average decrease in accuracy caused by removing one variable from the set of predictors. Variables included in the model can thus be ordered according to their contribution to model performance.

### 2.6 Cross-validation, model fitting and evaluation

Imbalanced dataset (49010 pseudo-absences vs. 4200 presences, Figure 2, panels A-B) are known to affect RF predictions (Khalilia et al., 2011). Therefore, it is recommended to use repeated random down-sampling, i.e. repeatedly fit the model on a subset of the original data containing all presence data and a random subsample of the same number of pseudo-absences (Japkowicz & Stephen, 2002; Valavi et al., 2022). As we subsample 7% of pseudo-absences among a globally distributed dataset, which displays spatial and underlying environmental structure (Figure 2, panels A-B), this structure is likely to be reflected in the output of random sampling and affect model predictions. Furthermore, model evaluation using standard cross-validation based on random sampling of observations is known to perform poorly when applied to ecological data with spatial or temporal structure (Ploton et al., 2020; Roberts et al., 2017). Thus, we split our dataset into blocks based on environmental dissimilarity among observations (Figure 2, panels C-D) and used this structure both in down-sampling and in an environmental blocks cross-validation strategy (Roberts et al., 2017; Valavi et al., 2019). Using the environmental blocks structure in down-sampling enforces sampling of pseudo-absences in diverse environmental groups while ensuring equal representation with presence data. In the cross-validation strategy, it allows to increase independence of validation data (held-out from training) and improves predictions error estimates (Ploton et al., 2020; Roberts et al., 2017).

The construction of the environmental blocks is described in detail in Supplementary Information 3 and eventually resulted in the definition of 7 environmental blocks (Figure 2, panels C-D).

We repeatedly fit 100 RF models using the randomForest R package 4.6-14 (Liaw & Wiener, 2002) to predict the probability of any observation to be a presence record, using environmental variables as predictors (number of trees = 1000). At each repetition of model fitting, we applied a 7-fold cross-validation strategy using the environmental blocks as folds, i.e., one block is held out, while the model is trained on the remaining 6 blocks. Within each block we sampled an approximatively equal number of pseudo-absences than the number of presences in the block, whereas pseudo-absences are sampled using a weighting scheme that prioritize observations that are further apart from presences according to the environmental dissimilarity (see Supplementary Information 3). These precautions ensure that data are balanced both within and between blocks, and that pseudo-absences (for which we have less confidence) are sampled at a certain environmental distance from presence data. Raster cells associated with the Azores archipelago (131 cells), which all reflect presence data, were systematically included in the training set, and never used for validation. At each repetition and for each 7-fold cross validation, we retrieved i) the variable importance as returned by the RF algorithm; ii) probabilistic predictions on all raster cells corresponding to landmasses of the Northern Hemisphere (N=8148045 cells); and iii) the three following model performance metrics computed on the held-out fold:

- sensitivity, which measures classification capacity, i.e. the ability to correctly classify occupied sites as suitable, based on the majority vote predictions of RF model (Fielding & Bell, 1997);
- Area Under the ROC Curve (AUC), which measures discrimination capacity, i.e. the ability to separate between occupied and unoccupied sites, regardless of any threshold value, based on the probabilistic prediction of RF model (Sillero et al., 2021);
- Continuous Boyce Index (CBI), a metric designed for evaluating SDM with presence-only data, which reflects the agreement between predicted probabilities of occurrence and observed proportions of sites occupied. This index ranges between −1 and 1, representing bad and good predictions respectively, and 0 corresponding to a random predictor (Boyce et al., 2002; Hirzel et al., 2006).

### 2.7 Suitability maps

The suitability maps display the median of models’ predictions, calculated over the 100 repetitions, on the landmasses of Northern Hemisphere. Our maps focus on the Northern Hemisphere since it is the only hemisphere where the Japanese beetle has ever been recorded and it is less likely for the Southern Hemisphere to be reached. Indeed, there are biological constraints limiting the possibility of a viable population reaching the Southern Hemisphere: egg-laying adults only emerge in the peak of Northern summer, and even if accidentally introduced to the Southern Hemisphere, they would arrive during the Southern winter (Allsopp, 1996).

As mentioned earlier, predictions correspond to a probability of Pj presence, thus ranging in the interval [0,1]. Following Hirzel *et al*. (2006), we used the plot of Boyce Predicted to Expected ratio against suitability (P/E curves) generated for each model repetition, to transform the continuous [0,1] probability interval into a discrete number of classes. These classes should reflect model calibration and more objectively represent the levels of habitat suitability. This method, based on the shape and confidence interval around the curves, provides a more honest discretization of the unit interval than an arbitrary choice of thresholds (Arshad et al., 2012; Hirzel et al., 2006; Lee-Yaw et al., 2022; Sattler et al., 2007; Vaughan & Ormerod, 2005), see Figure 1 in Supplementary Information 4 for more details).

We used an independent test dataset, set aside from model fitting and performance evaluation, to provide an unbiased evaluation of the suitability maps (median of models’ predictions). This test dataset includes all Japanese beetle presence data in Continental Europe, as well as presence data recorded in the years 2020-2021 in North America, Japan and the Azores (4098 presence raster cells). We contrasted the distribution of these presence data with predicted suitability values. Finally, we computed the Multivariate Environmental Similarity Surface (MESS, Elith et al., 2010) to identify regions of the Northern Hemisphere where model predictions constitute an extrapolation due to greater difference between environmental factors there and those used in model training (Figure 3A & Figure 2 in Supplementary Information 4).

**Figure 3 :**
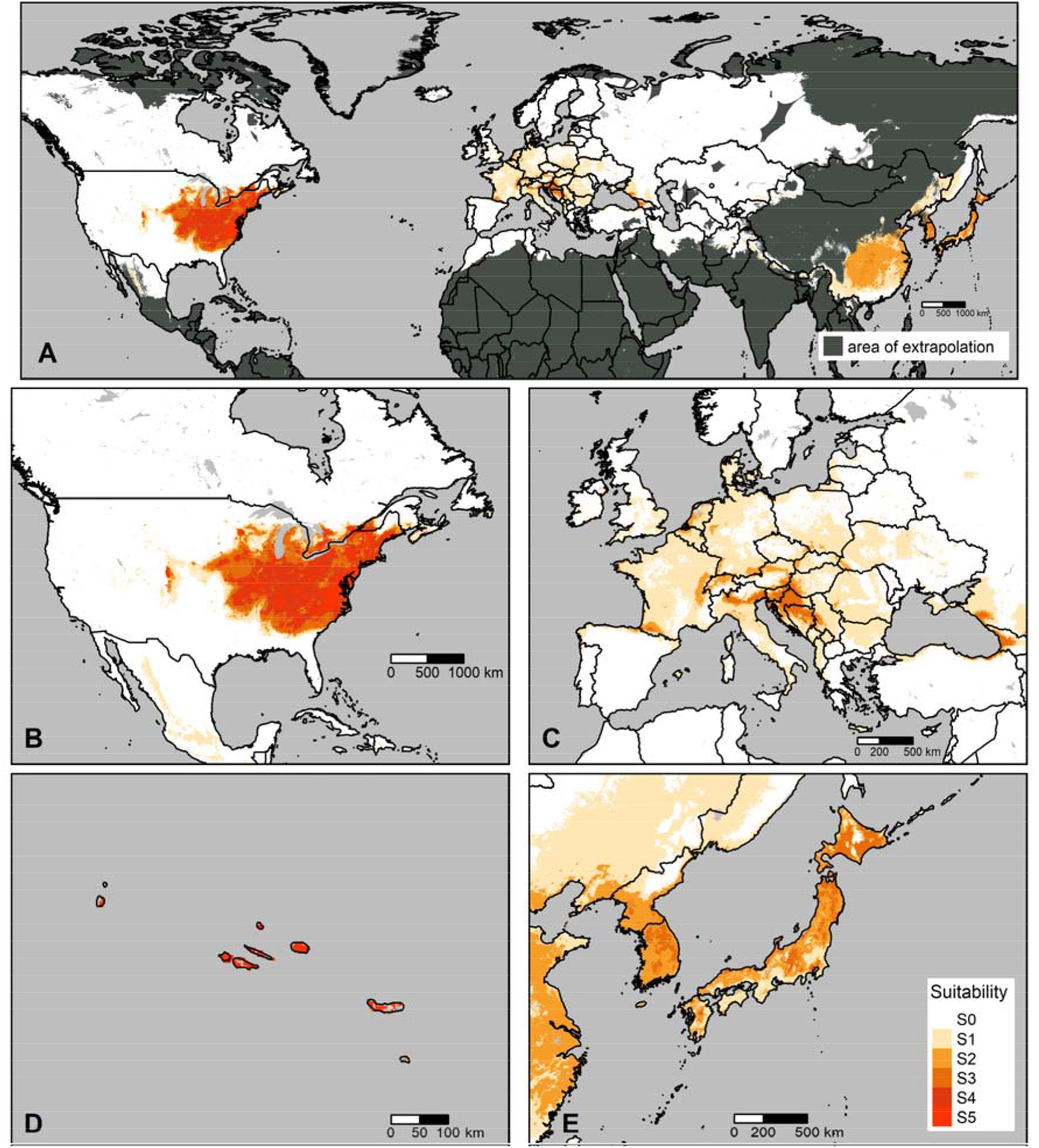
Suitability maps for the Japanese beetle. A) Northern Hemisphere, displaying areas of extrapolation in dark grey (Supplementary Information 4), B) North America, C) Continental Europe, D) Azores archipelago, E) Japan. Scales vary among panels and are indicated in kilometers on the maps. Darker colors correspond to greater suitability for the Japanese beetle.

## 3 Results

### 3.1 Model performance and calibration

All measures of model performance point towards a well-calibrated model with consistent and robust outputs throughout all 100 repetitions. The AUC varies between 0.72 to 0.98 with a mean 0.88 (SD=0.07) reflecting a good discrimination capacity, while sensitivity ranges between 0.33 and 1 with a mean 0.82 (SD=0.2), reflecting a good classification capacity. Furthermore, the Continuous Boyce Index (CBI) varies from 0.93 to 1 with a mean of 0.99 (SD=0.007), reflecting a very good calibration of the models.

The P/E curves method allowed to identify six classes of suitability that best reflect model calibration. Suitability maps and related results will thus be presented using these classes of increasing suitability. The first class, hereafter referred as S0, corresponds to suitability values between 0.00 and 0.19 and indicates an environment that is least favourable to the Japanese beetle (i.e. 97.5% of models predict less presences than expected by chance). For the following class S1 = [0.19 to 0.35], between 2.5% and 50% of models predict more presences than expected by chance. Class S2 = [0.35 to 0.45] reflects an ability between 50% and 97.5% of models to predict more presences than expected by chance. For these two classes, the presence of the beetle cannot be excluded, but uncertainty remains about long-term suitability for the organism Finally, the remaining classes indicate an increasing level of suitability for the beetle where at least 97.5% of models predict more presences than expected by chance (S3 = [0.45 – 0.67]), more than half of models predict at least 5 times more presences than expected by chance (S4 = [0.67 – 0.84]) and more than half of models predict at least 10 times more presences than expected by chance (S5 = [0.84 – 1.00]).

Overall, the proportion of presences from the test dataset increases along classes of increasing predicted suitability, and 93% of presences from the independent dataset are located within cells of the last three classes of suitability (13.5% in S3, 27.5% in S4 and 52% in S5, respectively). Only 0.6% of test presences are located within cells of class S0.

### 3.2 Suitability maps

Model predictions are shown in Figure 3 for the Northern Hemisphere with a focus on North America, Continental Europe, Japan and the Azores archipelago. A great proportion of northeastern USA and adjacent areas of southeastern Canada are predicted as highly suitable. Another cluster of high suitability is found at the foothills of the Front Range, east of the Rocky Mountains and is connected with the eastern high-suitability cluster via a corridor of moderate suitability crossing the Great Plains. In Mexico, the elevations of the Sierra Madre Occidental Mountain range constitute another continuous patch of suitable land. Smaller and scattered patches of low suitability may be found west of the Rockies. In the rest of the North American continent, it prevails the S0 class. In Japan, the native range of the beetle, we observe a gradient of moderate (S1) to high (S2 and S3) suitability towards the northern part of the archipelago. South Korea and eastern China were also predicted to be moderately suitable. The Azores islands are predicted to be highly suitable (mostly S4 and S5). Central Europe is almost entirely suitable (being at least class S1), with a continuous cluster of higher suitability (S2 and S3) stretching from the foothills of Western Italian Alps to the northern Balkans and the western Pannonian basin. Another cluster of higher suitability is found along the southeastern shores of the Black Sea. Smaller and scattered patches of moderate suitability can be found in the northern foothills of the Alps (France, Germany, Austria, Switzerland and Slovenia), west of the Pyrenees and coastal Low Lands (Belgium and the Netherlands).

The repartition of cells by suitability classes is shown by major geographical areas in Figure 4. In Japan, the two highest classes of suitability (S4 and S5) are not observed, while 3% of cells are considered as least suitable (S0). In North America (Canada and USA) we observe a rather sharp classification between suitable and least-suitable areas, with more than 80% of the domain considered as least suitable (S0), and the remaining cells distributed quite evenly across increasing suitability classes (Figure 4). The Azores archipelago is almost entirely highly suitable with 93% of the cells distributed in the highest suitability class S5. In Continental Europe, 62.7% of the domain is considered as least suitable (S0), 32.7% of the domain falls within the S1 class, 3.2% into class S2 and 1.4% in class S3. Classes of higher suitability (S4 and S5) are not observed although.

**Figure 4 :**
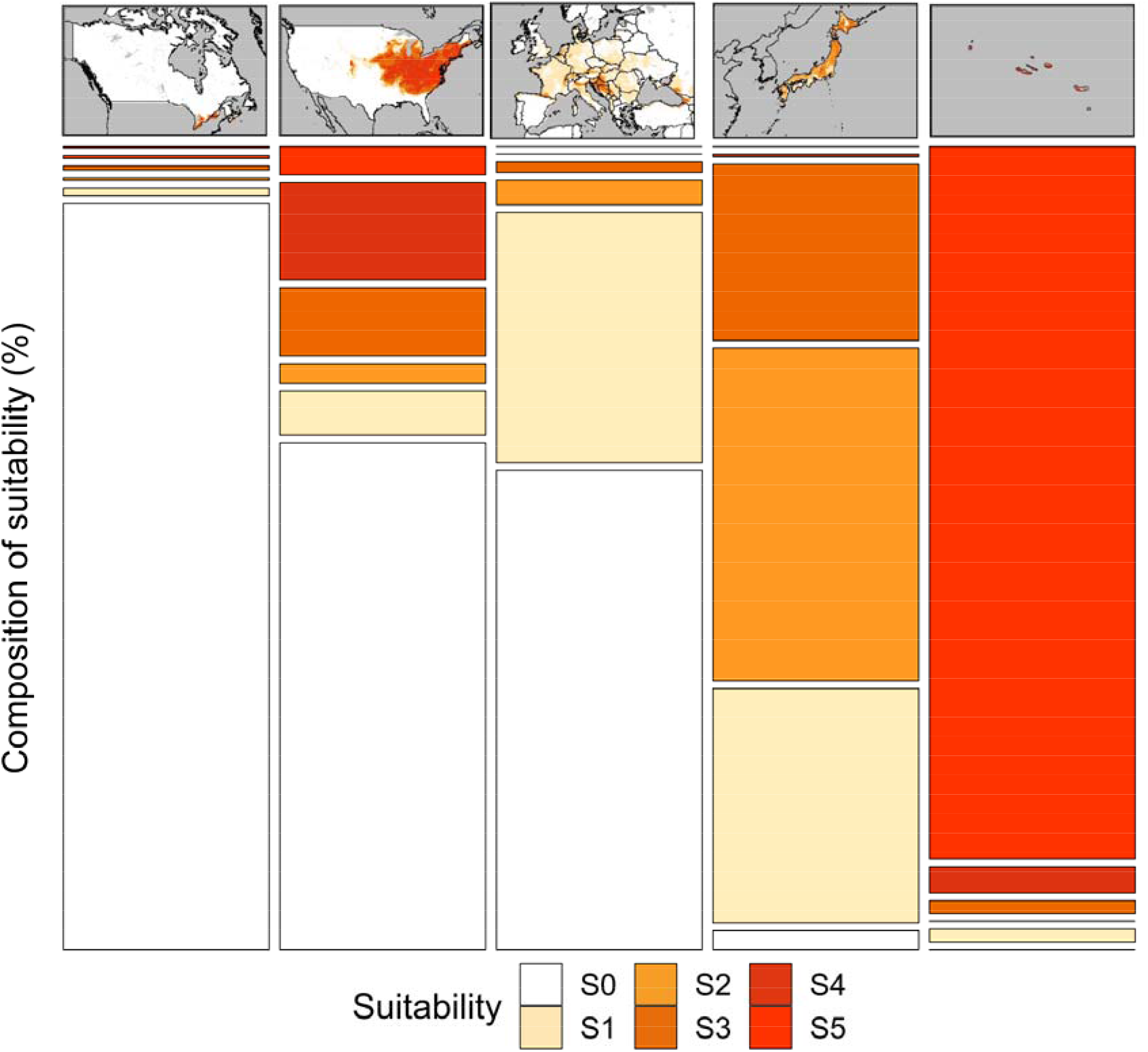
Composition of suitability, i.e. the proportion of total area covered by each suitability class, in Canada, USA, Continental Europe, Japan and the Azores archipelago. Darker colors correspond to greater suitability for the Japanese beetle.

### 3.3 Variable importance

The variable importance associated with each of the 133 model predictors is shown in Figure 1 in Supplementary Information 5. Clearly, the variable importance is not evenly distributed across variables: 50% (resp. 80%) of the cumulative importance is captured by the first 26 (resp. 57) most important variables. Three predictor themes contribute most to model predictions: climatic, human-related and land use predictors. These three predictor themes are similar in terms of importance as variable importance does not split predictors thematically. Regarding climatic predictors, we find average, minimum and maximum air temperatures, average soil temperature (between 5 cm and 15 cm), precipitations, growing degree days, actual evapotranspiration, water vapor pressure, downward surface shortwave radiation and runoff amongst the 26 predictors accounting for 50% of cumulative importance. These important climatic predictors are either annual average values, seasonal average values (implicitly representing seasonality), or some type of standard deviation of values (explicitly representing seasonality). Most of the important climatic predictors are implicitly or explicitly reflecting seasonality. Predictors which are explicitly reflecting seasonality are isothermality, temperature seasonality, annual mean diurnal range, precipitation seasonality. Predictors which are implicitly reflecting seasonality are summer and fall actual evapotranspiration, water vapor pressure, minimum and maximum temperatures, precipitations; precipitation of Warmest Quarter, and spring soil temperature between 5 cm and 15 cm.

The second group of important predictors are human-related variables. Human population density, travel time to the nearest cities and density of roads are amongst the predictors accounting for 50% of cumulative importance. Distance to the closest airport completes this list when considering the predictors accounting for 80% of cumulative importance.

Some land-use predictors also contribute to the cumulative importance: deciduous broadleaved trees patches of forest close to open area (“Tree cover, broadleaved, deciduous, closed to open (>15%)”) and urban areas.

## 4 Discussion

The main objective of this study was to provide robust and accountable predictions of the Japanese beetle suitability range in Europe, a fundamental step in order to design appropriate surveillance, containment and eradication strategies. We applied a machine-learning approach to study Pj distribution based on past observations of the beetle presence in its native and long-infested range, producing an accurate model for predicting its suitability in the Northern Hemisphere, with a particular focus on the European continent. There, our predictions highlight the presence of clusters of high suitability, embedded in an overall moderate suitability matrix, most of which are still not currently infested. This confirms that Pj invasion of Europe could spread further from its current distribution with potential high pest impact, reinforcing the need for a context-specific risk-based surveillance strategy.

### 4.1 Species distribution modelling

#### 4.1.1 SDM of invasive species

A critical point for the training of SDM is to constitute a database that is well representative of the environmental conditions in regions where the organism is present, as well as where it is absent, allowing the model to neatly separate those environments that are favourable from those that are not. Since we are modelling the distribution of an invasive species, we excluded recently infested regions (i.e., Continental Europe) from model training, as species distribution in these regions mostly reflects dispersal capacities rather than conditions which are favourable to the species. Although previous authors did include data from Continental Europe in model training (Della Rocca & Milanesi, 2022; Kistner-Thomas, 2019), our decision not to include them is supported by recent ecological literature (Liu et al., 2020), which proves that with reduced residence time, the invaded niche is smaller than the native niche with large unfillings, i.e. favourable conditions in the native niche are not occupied in the invaded niche.

#### 4.1.2 Citizen science data, sampling bias and target-group strategy for pseudo-absences

In the lack of a standardized and comprehensive database with Pj presences at global scale, we used opportunistic citizen-science observations to constitute the presence-only database, which, on one hand, ensured a global coverage with tens of thousands of presence records, but that, on the other hand, may affect the quality of the results (Dobson et al., 2020; Steen et al., 2019). In particular, opportunistic citizen-science data are likely to be geographically biased towards easily-accessible locations, typically urban, agricultural and recreational areas (August et al., 2020). To compensate for the sampling bias and lack of absence data, we chose to generate pseudo-absence data using the target-group strategy (Phillips *et al*., 2009). This approach has been confirmed to be unaffected by heterogeneous sampling effort, if the target group includes generalist species and occurrences in the widest environmental subspace associated with the study domain (Botella et al., 2020). We hence chose *Coleoptera* as our target-group because it is the most diverse group of insects and includes many generalist species (Bouchard et al., 2017; Lövei & Sunderland, 1996). By comparing the performance of random vs. target-group strategy in terms of variable importance (not shown here), we noticed a dramatic reduction of the relative importance of all human-related variables such as population density, urban land cover and roads densities. In terms of suitability maps, the random-sampling strategy produced a patchy distribution of high-suitability pockets in correspondence of big cities surrounded by low suitability areas, while the target-group approach produced a more evenly-distributed suitability envelope with varying levels of suitability, which is more likely representative of the patterns driven by underlying biological and ecological processes. Both results confirmed the presence of spatial bias in the data, whose effects have been mitigated by using the target-group approach. It is worth noticing that human-related factors have not disappeared from the list of important variables. This could account for the fact that the Japanese beetle has a certain preference for anthropogenic environments due to the presence of favoured food sources and irrigated turfs suitable for oviposition (Althoff & Rice, 2022; Fleming, 1972), as well as reflect its hitchhiking behaviour. It is challenging to disentangle what proportion of the relative importance attributed to human-related variables is actually explained by the beetle preference and what should be attributed to remaining sampling bias: more research needs to be conducted on this issue.

#### 4.1.3 Variable importance

The machine learning approach provides a ranking of the predictors in terms of relative importance: we found that climatic and human-related variables are most relevant for the beetle’s distribution, in agreement with previous works (Della Rocca & Milanesi, 2022; Kistner-Thomas, 2019; Zhu et al., 2017). In particular, we showed the importance of climatic seasonality, with sites displaying higher annual variation and lower daily variation being more favourable. This result reinforces the belief that Pj is well adapted to temperate climate. Our results also highlight that variables such as actual evapo-transpiration and vapor pressure are very important and we believe that they could be proxy of water availability in soil, related to favourability of soils for egg-laying by adults and larval survival. Similarly, the most relevant land-use variable is associated to broadleaved forest in proximity to open spaces, which corresponds to a landscape where two food resources (broadleaved trees and open spaces, typically crop fields) are in close proximity, also providing shaded areas along the boundaries that can preserve soil humidity during summer and are hence favourable to oviposition.

### 4.2 Suitability predictions in long-infested areas

In Japan, there are few observations available from the citizen-science platform, so it is difficult to evaluate the quality of the predictions. Nonetheless, most of the Japanese archipelago (97%) is classed within the suitability classes S1, S2 and S3. Interestingly, very few areas are classed at the lowest and highest levels of suitability (2.5% in S0, 0.3% in S4 and none in S5, respectively). This could be explained by a difference in crop types between Pj’s native and invaded niches, and the presence in Japan of natural predators and parasites, which regulate Pj populations (Clausen et al., 1927). In the Azores archipelago, where the beetle has been introduced in the seventies, the suitability predicted by the model is always very high (majority of classes S3, S4 and S5). The invasion history and the difficulties in eradication programs reported by local authorities support this prediction (Vieira, 2008).

The situation in North America, especially in the USA, is more paradigmatic and deserves a deeper discussion. The first report of the beetle dates back to 1916 in a New Jersey nursery (Dickerson & Weiss, 1918) and the beetle’s distribution extended westward. Pj is currently established in 28 US states, and at least 13 more US states and Canadian provinces have reported interceptions, likely consequence of human-mediated transport (Althoff & Rice, 2022; CERIS, 2020; USDA-APHIS 2020). In 1998, the USA established the U.S. Domestic Japanese Beetle Harmonization Plan, lately revised in 2016 (NPB, 2016). This plan assigns each US state to one of four categories, according to the beetle infestation status, risk of entry via artificial means and natural spread, expected impact established by a pest risk assessment, and the need for quarantine certification protocols. To further evaluate our suitability maps, we use the NPB’s categorization (2016) as a reference for Pj current distribution in the USA. However, although we included factors such as distance to the closest airport and density of roads in our model, our maps do not reflect the risk of entry per se, nor do we consider the potential pest impact (damage on crops, turfs, etc.).

We predicted lowest suitability (S0) for Florida and Louisiana (Figure 3B), which is in agreement with their classification as Category 4 states, “*Historically Not Known to Be Infested/Unlikely to Become Established*” (NPB, 2016). We predicted mostly lowest suitability with localized clusters of higher suitability (class S3) in Wyoming, also a Category 4 state, which is supported by recent interceptions of Pj (Althoff & Rice, 2022). We predicted the Western US states, classified as Category 1- “*Uninfested/Quarantine Pest*”, as least-suitable (S0) with very small and disconnected patches of low suitability (S1) (Figure 3 and Table 1). However, Category 1 indicates that official pest risk assessments found a moderate to high pest impact in case of establishment that can be mitigated by applying quarantine protocols and eradication programs. This discrepancy can be explained by the presence of high-value crops such as grapes and berries, that are amongst the preferred hosts for the Japanese beetle (Althoff & Rice, 2022) and significant risk of accidental human-mediated introduction in these states due to the presence of highly-connected hubs for commercial and passenger flows. Successful eradications of past introductions in California and Oregon support our predictions of low suitability (Althoff & Rice, 2022). Comparison with Category 2 (“*Uninfested or Partially Infested”)* is tricky as it includes both uninfested and partially infested states, where NPB (2016) found that Pj could survive but pest impact would be low to moderate. In these states, we mainly predicted lowest suitability (82% of cells in class S0) and some clusters of high suitability (6% in classes S2 or higher). For example, Colorado, which we predicted to have a high-suitability cluster in the Eastern foothills of the Front Range, and Texas, which we predicted as mostly S0, are both assigned to Category 2 according to NPB (2016). We believe our findings are mainly in agreement with this classification, as even if small local populations may be sustained in managed lands (turfs and lawns for example), the greatest proportion of states is least suitable to the beetle. Finally, in Eastern USA, Category 3 - “*Partially or Generally Infested”*, corresponds mostly to highest levels of suitability (73% classed as S2 or higher) (Table 1). Overall, we conclude that our suitability predictions agree with the official classification of NPB (2016) and that observable differences ascribe to the fact that we did not explicitly account for risk of entry and pest impact.

**Table 1 :**
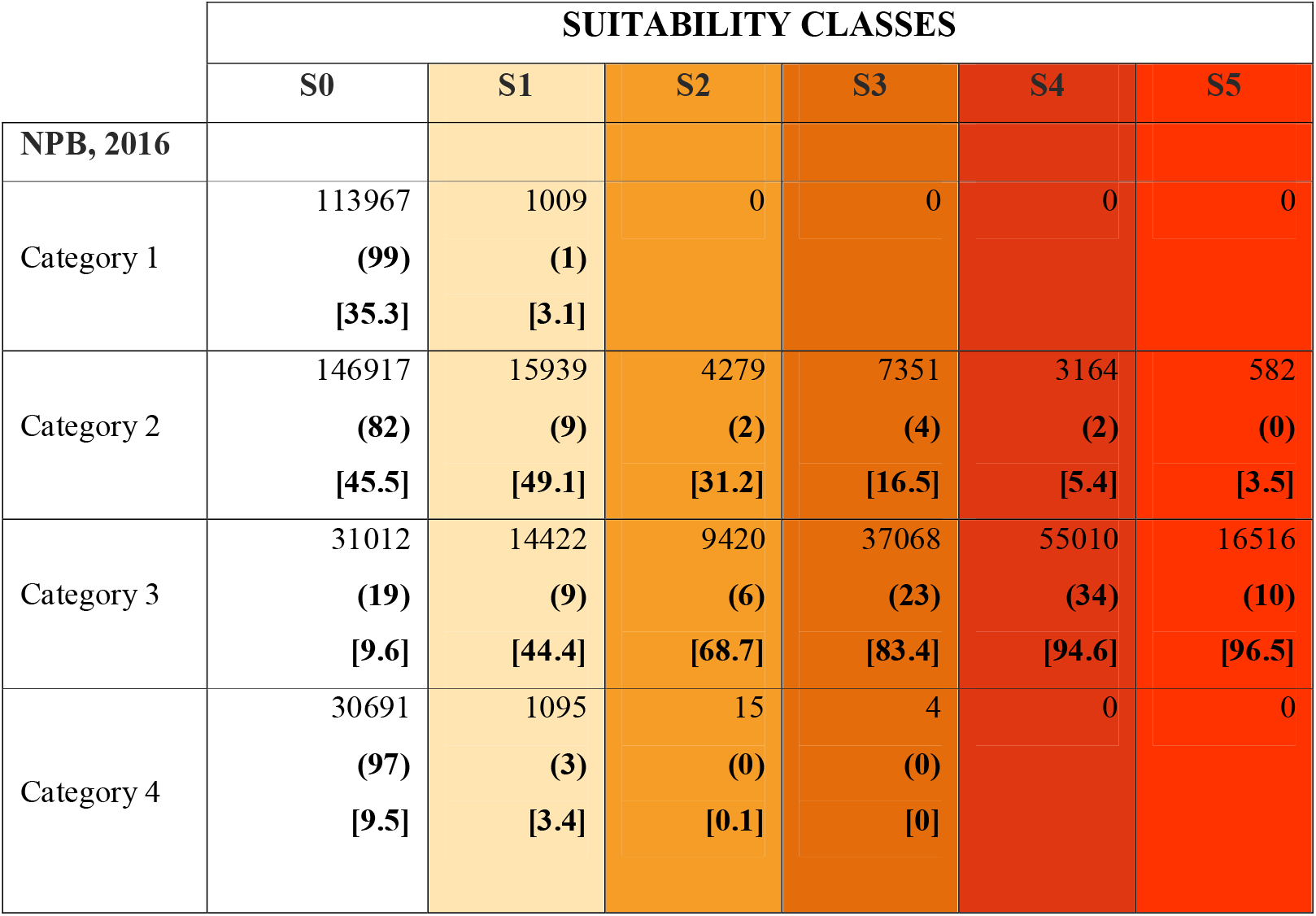
Distribution of predicted suitability in the USA among the categories of Pj infestation levels reported by the NPB (2016). Values indicate the number of cells in each suitability class and category; values in parentheses represent the percentage w.r.t. the corresponding row of the table; values in brackets represent the percentage w.r.t the corresponding column of the table. USA categories according to NPB (2016): Category 1 = “Uninfested/Quarantine Pest”, Category 2 = “Uninfested or Partially Infested”, Category 3 = “Partially or Generally Infested” and Category 4 = “Historically not known to be infested / Unlikely to Become Established”.

Recent proposals by Zhu et al. (2017), Kistner-Thomas (2019) and Della Rocca & Milanesi (2022) also produced suitability maps for the Japanese beetle in the USA, and major differences can be observed. First of all, both Zhu et al. (2017) and Kistner-Thomas (2019) predicted, with varying degrees of suitability, the entire West Coast as suitable, with multiple narrow “corridors” from the infested Central and Eastern US towards the West Coast, through the Rocky Mountains and Sierra Nevada. Della Rocca & Milanesi (2022) predicted a cluster of high suitability in the northern Pacific Coast (Washington and Oregon states) and low-to-moderate suitability along the Southern Rockies. These results are in conflict with our predictions and Allsopp (1996) statement, who affirmed the unlikelihood of westward expansion due to unfavourable conditions in the Rocky Mountains and Sierra Nevada. Even more striking is the difference with respect to Southern and Northern US states and Canadian provinces: previous authors predicted these states as highly suitable, at least as suitable as the most infested areas in Northeastern US. This is in strong contradiction with our results, as well as the classification by (NPB (National Plant Board), 2016). The fact that Pj has not established yet in these regions, despite the fact that they had decades to reach them from northern USA (both by natural and human-assisted spread) and that reduced local populations have already been observed there, without ever becoming established further support our results (Althoff & Rice, 2022; CDFA, 2022; Koppenhöfer et al., 2000).

In the light of these considerations, we can confirm that our model is capable of producing reliable predictions about habitat suitability for the Japanese beetle, and that compared to previous proposal in the literature, it is less prone to overestimation (i.e. tendency towards false positive, labelling as suitable those areas that apparently are not). This could partly be attributed to our use of an environmental-blocks approach for pseudo-absence selection in model fitting (Valavi et al., 2019). Based on this last remark, we can now focus on the European continent, where an infestation is ongoing and the need for reliable suitability maps are urgent.

### 4.3 Projection of suitability in Continental Europe and comparison with previous studies

Our predictions depict a complex scenario for Europe (Figure 3C), with two main clusters of high predicted suitability: the first in the foothills of the Alps (both North and South) and in the Northern Balkans, and the second on the eastern shores of the Black Sea. It is worth noticing that the present distribution of the Japanese beetle in Europe (Piedmont and Lombardy regions in Italy, Ticino canton in Switzerland) is concentrated in areas that we predict as the highest suitable in the continent. Smaller and patchier clusters of moderate suitability can be found in Southwestern France, close to the Pyrenees, in Brittany, and in the Low Lands (coastal Belgium and Netherlands). One major issue is that a great proportion of Central Europe is considered as suitable, even if at the lowest level marginal suitability (class S1), and that all clusters of moderate and high suitability are embedded into this low-suitability area, creating a continuum of suitable land across the continent. The areas predicted as least-suitable are found along the coastline of the Mediterranean Sea (Iberic Peninsula, Southern Italy, Greece, Anatolia), in the northern part of the continent (British islands, Scandinavia, Baltic rim) and Eastern Europe. Even if the general picture is overall quite worrying, especially for Central Europe, it bears no comparison with the predictions from previous works who considered the entire continent as highly-suitable (Della Rocca & Milanesi, 2022; Kistner-Thomas, 2019; Zhu et al., 2017), with the only exception of the Iberic Peninsula, the peaks of the Alps and Scandinavia. As expected, those models that were overestimating the suitability in the training dataset (mostly North America), maintained that tendency when extrapolating to Europe. Our maps highlight a more contrasted suitability scenario for Europe, which is an improvement for prioritization in surveillance.

By looking at how our suitability classes distribute among the categories of NPB (2016) (Table 1), we could try to infer the expected infestation levels and pest-impact in Europe. Areas classified at S3 and S4 levels are very likely to observe the same levels of infestation as the Eastern part of USA (83% and 69% in Category 3 resp., Table 1), with high damage levels to crops and continuous occupations. Areas with the lowest level of suitability (S0) are very likely to display low infestation and low pest-impact (45.5% in Category 2). On the other hand, cells at suitability level S1 bear more uncertainty as they split uniformly between Category 2, where Pj could survive but pest-impact is moderate to low, and Category 3, considered generally infested, where the “spread cannot be effectively slowed, and regulation of host commodities is not likely to be effective” (NPB, 2016). Therefore, the class S1 is the less informative, as it could reflect infestation levels as in New Jersey (overall highest suitability levels), Colorado (clusters of very high suitability in overall moderate suitability) or Texas (overall lowest suitability). Although the USA invasion is very informative, transferring the NPB (2016) categorization to Europe to infer infestation levels has its limits, since the European and American landscapes and policies are quite different. Furthermore, a niche shift cannot be excluded in Europe as the invasion is still recent and terrestrial ectotherms are known to show higher level of niche expansions (Liu et al., 2020). Besides, Liu et al. (2020) found that introductions from Asia to North America and from North America to Europe show a lower level of niche conservatism. Future research could help discriminate this uncertainty with updated and long-term data on the European invasion. Most importantly, analysis which would incorporate risk of entry and potential damage could bring information to further discriminate the importance of these cells for Pj invasion in Europe.

### 4.4 Implications for controlling the ongoing invasion of the Japanese beetle in Europe

Building a surveillance strategy for Europe is crucial and mapping the environmental suitability is a major milestone. However, several pieces of information still need to be added to get the full picture: connectivity to the infested zones, accounting for both natural and human-mediated pest dispersal, combined with spatial cost assessment and environmental suitability, could provide a rationale for selecting surveillance sites and proportionate eradication strategies. Since environmental suitability is heterogeneously distributed across the continent, and the connectivity network of Europe transports imposes a non-trivial geometry for the introduction of the beetle into new areas (Brockmann and Helbing, 2013), we call for a mixed surveillance approach: in already infested areas, efforts should focus on the containment of growing populations and tracking the front of invasion in neighbouring non-infested areas. Furthermore, quarantine certification protocols should be applied to regulated products from infested area (plants and plant material, such as soil, compost, mulch, sod, etc.), to ensure beetle-free shipments especially during the adults’ flight period (June to September). Similarly, transportation hubs located within the infested area should be inspected and all destinations directly reachable from there should surveil (visual inspections and traps in the surroundings of the hubs) and be ready to react to the potential arrival of the beetle. The quarantine protocols and eradication efforts applied in areas with potential high pest-impact in Western USA demonstrated that timely reaction can be crucial for controlling the spread of the beetle (CDFA, 2022). Efficient and rapid exchange of information, data and knowledge between phytosanitary agencies of EU regions and countries, are key to prevent unnecessary delays in the reaction. Coordinated effort should be implemented from EU agencies. Finally, citizens and stakeholders should be sensitized via appropriate media campaign to seek for their engagement in the detection and fight against the invasive Japanese beetle.

## Author Contribution Statement

S.P. and D.M. conceived the initial project. S.P. acquired research funding. L.B., S.P. and D.M. discussed the data and methodology requirements for the study. L.B. performed data collection, implementation of the computer scripts, data visualization and analysis. L.B., D.M. and S.P. analyzed data and results. L.B., S.P. and D.M. wrote, reviewed and edited the manuscript. All authors reviewed the results and approved the final version of the manuscript.

## Key Message

- Suitable areas for the Japanese beetle are heterogeneously distributed across Europe
- Central Europe is suitable and the area currently occupied is among the most suitable in Europe
- Only 1% of the overall most suitable area at risk of establishment in Europe is currently occupied
- Seasonality, land use and human settlements greatly affect site suitability to the Japanese beetle
- Distribution of suitability in Europe calls for a mixed surveillance strategy

## Acknowledgements

This research was supported by the IPM-Popillia project, funded by the European Union Horizon 2020 research and innovation programme under grant agreement No 861852. Authors would like to thank Servizio Fitosanitario Ticino, Plant Health Service - Piedmont Region (Italy), Secretaria Regional da Agricultura e do Desenvolvimento Rural do Governo dos Açores, and the Canadian Food Inspection Agency, for providing standardized survey presence data included in analyses. Authors would also like to thank citizen science platforms and all citizens who take part in citizen science surveys, for their contribution to the advancement of research.

## 6 Statements and Declarations

### 6.4 Data Availability

The datasets generated during and/or analysed during the current study are available in the French Research Government repository, https://doi.org/10.57745/GM2YVL

