## Supplementary Information 1 for "A new chapter of the Japanese beetle invasion saga: predicting suitability from long-infested areas to inform surveillance strategies in Europe"

*Author names : Leyli Borner[1], Davide Martinetti[2], Sylvain Poggi[1].*

*Affiliations :*

*1. INRAE, Institut Agro, Univ Rennes, IGEPP, 35653, Le Rheu, France*

*2. INRAE, UR 546 BioSP, Avignon, France*

### Supplementary Information 1: Presence data sources and pre-processing

These data were extracted in November 2021 using the spocc 1.2.0 R package (Chamberlain et al., 2021) and the iNaturalist query tool. We removed unreliable identifications, preserved specimen from collections and data with imprecise coordinates using the scrubr 0.4.0 R package (Chamberlain, 2020). We removed duplicate observations, and temporal series of observations made at the same location within the same year. At the end of the data cleaning stage, the dataset contains 53678 geo-referenced presence data, of which 78% originate from citizen science platforms (see Figure 1 in the main text & Table 1 in this supplementary information). The earliest Japanese beetle observation originates from the iDigBio platform and is dated August 1917 in New Jersey (USA) (see Figure 1 in this supplementary information). The latest observations originate from iNaturalist and correspond to dead adults observed in November 2021 in Colorado and North Carolina (USA), and in Freiburg im Breisgau (Germany, interception).

All JB presence records used in this study are available on the French Research Government repository (Borner et al., 2022).

*Table 1 : Distribution of Japanese beetle presence data collected from years 2010 to 2019 among sources (N=25376)*

| Source | # of presence data | Type | Geographical extent |
|---|---|---|---|
| iNaturalist | 6328 | Citizen science | World |
| GBIF | 3800 | Citizen science | World |
| BISON | 3134 | Citizen science | USA & Canada |
| EDDMapS | 1871 | Citizen science | USA & Canada |
| iMapInvasives | 26 | Citizen science | USA |
| SCAN-Bugs | 20 | Citizen science | World |
| iDigBio | 9 | Citizen science | USA & Japan |
| FGF- Universidade dos Açores | 5146 | Stand. Surveys | Azores archipelago (Portugal) |
| Servizio Fitosanitario Regione Piemonte | 4979 | Stand. Surveys | Italy |
| Canadian Food Inspection Agency | 48 | Stand. Surveys | Canada |
| Servizio Fitosanitario Ticino | 15 | Stand. Surveys | Switzerland |

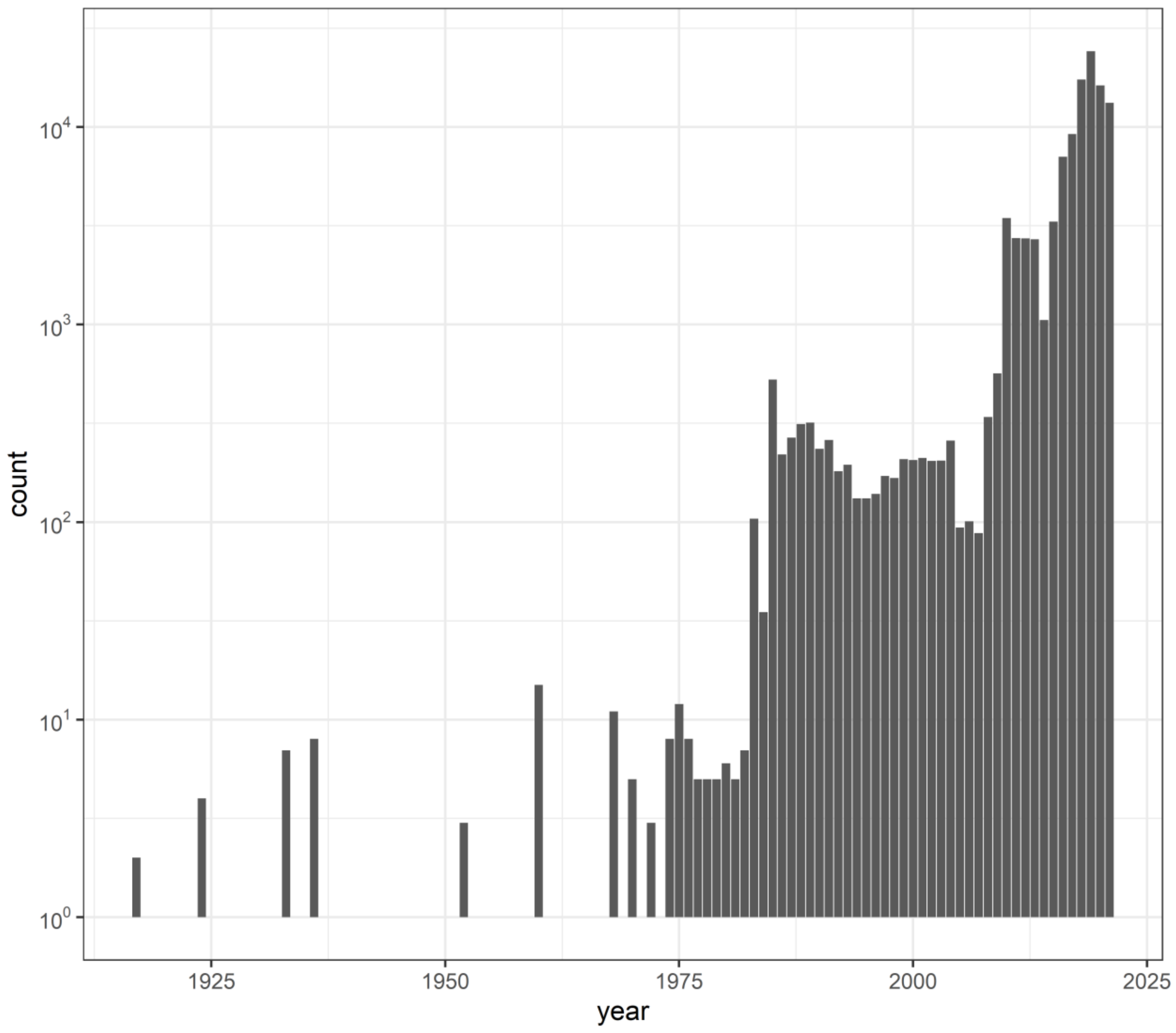


*Figure 1 : Number (log-scaled) of Japanese beetle presence data collected for this study across the years 1917 to 2021*
