## Supplementary Information 2 for "A new chapter of the Japanese beetle invasion saga: predicting suitability from long-infested areas to inform surveillance strategies in Europe"

*Author names : Leyli Borner[1], Davide Martinetti[2], Sylvain Poggi[1].*

*Affiliations :*

*1. INRAE, Institut Agro, Univ Rennes, IGEPP, 35653, Le Rheu, France*
*2. INRAE, UR 546 BioSP, Avignon, France*

### Supplementary Information 2 : Explanatory variables

All environmental predictors used in this study are available on the French Research Government repository https://doi.org/10.57745/GM2YVL (Borner et al., 2022).

*Table 1 : Explanatory variables used in modelling, source of the predictors, spatial and temporal resolutions, and short description of the 133 explanatory variables used in the model.*

| Source | Spatial resolution (in meters) | Temporal resolution | Short description |
|---|---|---|---|
| **OurAirports** | 20 | | large & medium airports; presence/absence of airports in 4km x 4km cells & distance from 4km x 4km cells' centroids to closest airport (https://ourairports.com/), time reference: accessed in September 2021. |
| **OpenLandMap** | 250 | average values | Soil characteristics variables; Clay content in % (kg / kg) ; Soil organic carbon content in × 5 g / kg (to convert to % divide by 2); Soil pH in H2O in × 10 ; Sand content in % (kg / kg); at 10 cm depth at 250 m resolution; To access and visualize maps use: OpenLandMap.org; time reference: 1950-2017 |
| **Soilgrids** | 250 | | World Reference Base (2006) Soil Groups; 30 classes; |
| **Climate Change Initiative (CCI) Land Cover V2** | 300 | | 23-class land cover map & Shannon index of diversity; layers available for 1998-2002, 2003-2007 and 2008-2012; time reference: 2008-2012 |
| **Composite Wetlands - Topography-Climate Index (CW-TCI)** | 500 | | Groundwater-driven wetlands from TCI (GDW-TCI(15%)), Regularly flooded wetlands (RFW), Intersection of RFWs and GDWs, Lakes (from HydroLAKES) ; Ardalan Tootchi, Anne Jost, Agnès Ducharne. Multi-source global wetland maps combining surface water imagery and groundwater constraints. Earth System Science Data, Copernicus Publications, 2019, 11, pp.189 - 220. 10.5194/essd-11-189-2019. hal-02045896 (https://hal.sorbonne-universite.fr/hal-02045896/file/essd-11-189-2019.pdf) |
| **Accessibility to cities** | 1000 | | Travel time to cities 2015 - The value of each pixel is the estimated travel time in minutes to the nearest urban area in 2015. Cities from population minimum:50,000 to population maximum: 50,000,000; time reference: 2015 |
| **CHELSA – Climatologies at high resolution for the Earth land surface areas. Version 1.2** | 1000 | annual average | Bioclim variables from CHELSA (http://chelsa-climate.org/) is a high resolution (30 arc sec, ~1 km) climate data set for the earth land surface areas. It includes monthly and annual mean temperature and precipitation patterns for the time period 1979-2013. Time reference: 1979-2013 |

| | | | |
|---|---|---|---|
| **Global Soil Bioclimatic variables at 30 arc second resolution** | 1000 | monthly average values | Soil temperature layers were calculated by adding monthly soil temperature offsets to monthly air-temperature maps from CHELSA (date range 1979-2013) (Karger et al. 2017, Sci Data); 0-5 cm and 5-15 cm depth; https://zenodo.org/record/4558732#.Yh3-AN_jKUk; averaged for spring, summer, fall and winter; time reference: 1979-2013 |
| **Gridded Population of the World, Version 4 (GPWv4) - 2020** | 1000 | | Population density - documentation: http://sedac.ciesin.columbia.edu/data/collection/gpw-v4/documentation, time reference: 2020 |
| **Worldclim 2.1** | 1000 | | Elevation - SRTM elevation data available from WorldClim 2.1 |
| **Global Roads Inventory Project - GRIP - version 4** | 4000 | | roads density in meters per km2 per cell for primary roads and secondary roads (https://www.globio.info/download-grip-dataset) |
| **Terraclimate** | 4000 | monthly - 2015 to 2020 | Maximum temperature, minimum temperature, vapor pressure, precipitation accumulation, downward surface shortwave radiation, wind-speed; Runoff, Actual Evapotranspiration, Soil Moisture, Snow Water Equivalent, Palmer Drought Severity Index, Vapor pressure deficit ; available from 1958 to 2020, we used monthly averaged for the years 2015-2020; computed seasons average for spring, summer, fall and winter; Global degree days, lifecycle and earliest date of emergence calculated from monthly minimal and maximal temperature; (http://www.climatologylab.org/terraclimate.html); time reference: 2015-2020 |
