## Supplementary Information 3 for "A new chapter of the Japanese beetle invasion saga: predicting suitability from long-infested areas to inform surveillance strategies in Europe"

*Author names : Leyli Borner[1], Davide Martinetti[2], Sylvain Poggi[1].*

*Affiliations :*

*1. INRAE, Institut Agro, Univ Rennes, IGEPP, 35653, Le Rheu, France*

*2. INRAE, UR 546 BioSP, Avignon, France*

### Supplementary Information 3: Construction of environmental blocks

We built the set of environmental blocks based on the following procedure:

1. Apply a decomposition in principal components to the environmental dataset (133 variables) for all cells used in model fitting (53079 cells) using Principal Component Analysis prcomp function (R Core Team, 2021);
2. Retain the components that explain 90% of the total variance (57 Principal Components);
3. Compute the weighted pairwise distances between all cells projected into the new space, using Gower distance to obtain a dissimilarity matrix (daisy function of cluster 2.1.3 R package, Maechler *et al.*, 2022);
4. Identify a set of homogeneous clusters minimizing the total within-cluster variance using hierarchical clustering on dissimilarity values corresponding to presence cells (Ward's method, hclust function, R Core Team, 2021));
5. Assign each presence raster cell to its corresponding block, and each pseudo-absence cell to the block assigned to the closest presence record in terms of dissimilarity value.

Raster cells associated with the Azores archipelago were set aside during this procedure because they were systematically assigned to an independent cluster as their corresponding environmental conditions are very peculiar and could not be found anywhere else.

*Table 1 : Distribution of data used in model training among regions and environmental blocks*

| | Environmental block | | | | | | |
|---|---|---|---|---|---|---|---|
| | **1** | **2** | **3** | **4** | **5** | **6** | **7** |
| **Countries** | | | | | | | |
| Canada | 399 | 5604 | 100 | 0 | 0 | 843 | 1150 |
| Japan | 496 | 14 | 0 | 13 | 21 | 0 | 2 |
| United States | 4035 | 4739 | 6589 | 11940 | 9048 | 4894 | 3192 |

*Table 2 : Distribution of presence and pseudo-absence data among environmental blocks*

| | Environmental block | | | | | | |
|---|---|---|---|---|---|---|---|
| | **1** | **2** | **3** | **4** | **5** | **6** | **7** |
| **Pseudo-absences** | 4530 | 9518 | 6611 | 11367 | 8516 | 4999 | 3469 |
| **Presences** | 400 | 839 | 78 | 586 | 553 | 738 | 875 |
