## Supplementary Information 4 for "A new chapter of the Japanese beetle invasion saga: predicting suitability from long-infested areas to inform surveillance strategies in Europe"

*Author names : Leyli Borner[1], Davide Martinetti[2], Sylvain Poggi[1].*

*Affiliations :*

*1. INRAE, Institut Agro, Univ Rennes, IGEPP, 35653, Le Rheu, France*

*2. INRAE, UR 546 BioSP, Avignon, France*

### Supplementary Information 4. Suitability maps

#### 1.1 Construction of suitability classes

To be able to rank our suitability predictions on different sites, we measured calibration of models (Lee-Yaw et al., 2022). Calibration measures the agreement between predicted probabilities of occurrence and observed proportions of sites occupied (for example, if 40% of sites with predicted probabilities 0.4 are occupied).

We used the plot of Boyce Predicted to Expected ratio against suitability (P/E curves) generated for each model repetition, to transform the continuous [0,1] probability interval into a discrete number of classes (Hirzel *et al.*, 2006). These classes should reflect model calibration and more objectively represent the levels of habitat suitability. This method, based on the shape and confidence interval around the curves, provides a more honest discretization of the unit interval than an arbitrary choice of thresholds (Arshad et al., 2012; Hirzel et al., 2006; Lee-Yaw et al., 2022; Sattler et al., 2007; Vaughan & Ormerod, 2005), see Figure 1 in this supplementary information for more details).

The first class, corresponds to suitability values between 0.00 and 0.19, indicates an environment that is least favourable to the Japanese beetle (i.e. 97.5% of models predict less presences than expected by chance). For the following class [0.19 to 0.35], between 2.5% and 50% of models predict more presences than expected by chance. Class [0.35 to 0.45] reflects an ability between 50% and 97.5% of models to predict more presences than expected by chance. For these two classes, the presence of the beetle cannot be excluded, but uncertainty remains about long-term suitability for the organism Finally, the remaining classes indicate an increasing level of suitability for the beetle where at least 97.5% of models predict more presences than expected by chance [0.45 – 0.67], more than half of models predict at least 5 times more presences than expected by chance [0.67 – 0.84] and more than half of models predict at least 10 times more presences than expected by chance [0.84 – 1.00].

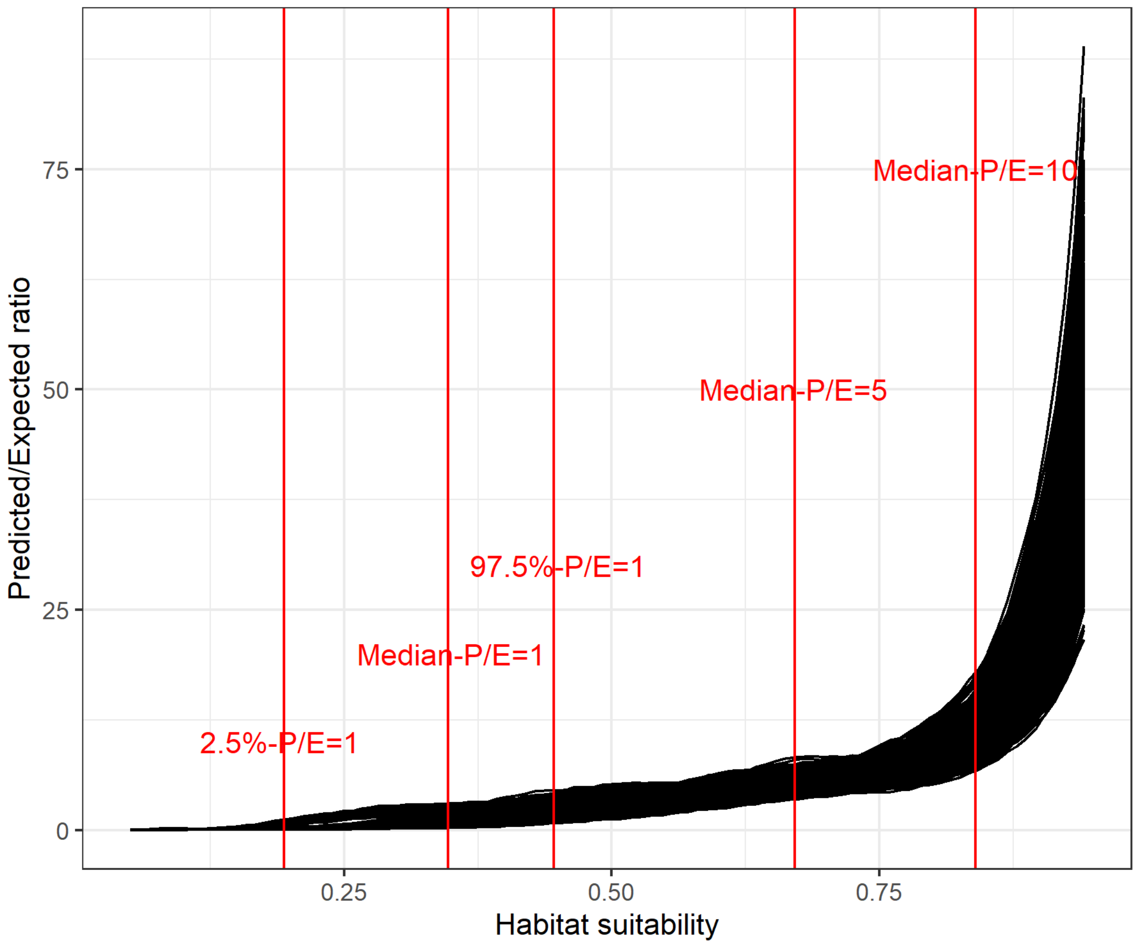


*Figure 1 : Boyce plot used to measure calibration and set thresholds to choose six suitability classes to map suitability*

##### 1.2 Global suitability maps and measure of extrapolation

In order to be able to identify areas of extrapolation in our predictions, we followed Elith et al., (2010) and computed the Multivariate Environmental Similarity Surface (MESS) using variables accounting for 80% of cumulated variable importance. In the main text we show a map of prediction and extrapolation on the Northern Hemisphere, here we provide the same information at the global scale. Figure 2 shows where model predictions constitute an extrapolation due to greater difference between environmental factors there and those used in model training. Some of our predictors are computed to reflect northern hemisphere seasons (e.g. Winter is from December to February and Summer is from June to August), which is why we found we are extrapolating on most of the southern hemisphere.

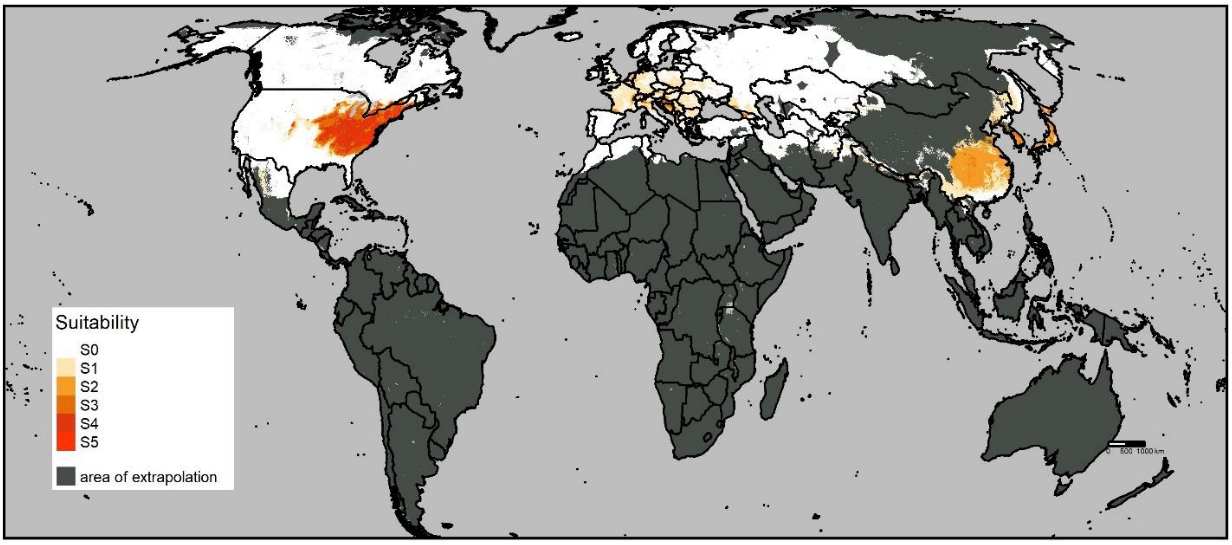


*Figure 2 : Global map of suitability and measure of extrapolation (computed with MESS using variables accounting for 80% of cumulated variable importance)*
