## Supplementary Information 5 for "A new chapter of the Japanese beetle invasion saga: predicting suitability from long-infested areas to inform surveillance strategies in Europe"

*Author names : Leyli Borner[1], Davide Martinetti[2], Sylvain Poggi[1].*

*Affiliations :*

*1. INRAE, Institut Agro, Univ Rennes, IGEPP, 35653, Le Rheu, France*

*2. INRAE, UR 546 BioSP, Avignon, France*

### Supplementary Information 5. Variable importance

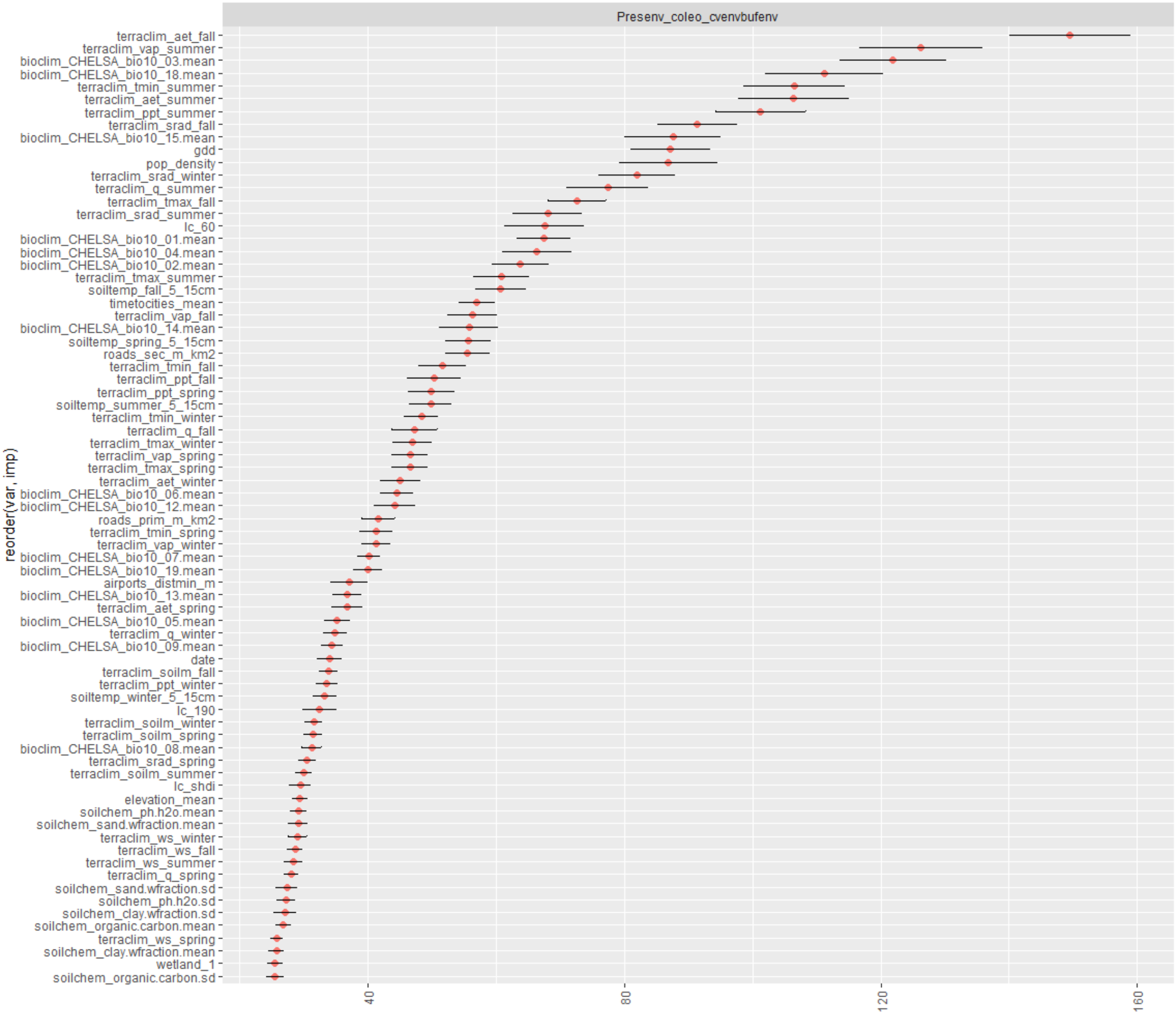


*Figure 1 : Mean and standard deviation of variable importance used in 100 repetitions of model training, 75 variables shown (for importance values superior to 25)*

*Table 1 : Importance of explanatory variables used in 100 repetitions of model training, 71 variables accounting for 90% of cumulative variable importance included*

| Position | Variable name | Mean Importance | SD Importance | Proportion of importance explained by variable | Cumulated importance explained by variables |
|---|---|---|---|---|---|
| 1 | terraclim_aet_fall | 149.5 | 9.5 | 3.6 | 3.6 |
| 2 | terraclim_vap_summer | 126.3 | 9.5 | 3.0 | 6.6 |
| 3 | bioclim_CHELSA_bio10_03.mean | 121.9 | 8.3 | 2.9 | 9.5 |
| 4 | bioclim_CHELSA_bio10_18.mean | 111.2 | 9.3 | 2.6 | 12.1 |
| 5 | terraclim_tmin_summer | 106.5 | 7.9 | 2.5 | 14.6 |
| 6 | terraclim_aet_summer | 106.4 | 8.7 | 2.5 | 17.2 |
| 7 | terraclim_ppt_summer | 101.2 | 7.0 | 2.4 | 19.6 |
| 8 | terraclim_srad_fall | 91.4 | 6.3 | 2.2 | 21.7 |
| 9 | bioclim_CHELSA_bio10_15.mean | 87.6 | 7.5 | 2.1 | 23.8 |
| 10 | gdd | 87.2 | 6.2 | 2.1 | 25.9 |
| 11 | pop_density | 86.9 | 7.7 | 2.1 | 28.0 |
| 12 | terraclim_srad_winter | 82.0 | 6.0 | 1.9 | 29.9 |
| 13 | terraclim_q_summer | 77.4 | 6.4 | 1.8 | 31.8 |
| 14 | terraclim_tmax_fall | 72.6 | 4.5 | 1.7 | 33.5 |
| 15 | terraclim_srad_summer | 68.0 | 5.4 | 1.6 | 35.1 |
| 16 | lc_60 | 67.5 | 6.2 | 1.6 | 36.7 |
| 17 | bioclim_CHELSA_bio10_01.mean | 67.5 | 4.2 | 1.6 | 38.3 |
| 18 | bioclim_CHELSA_bio10_04.mean | 66.4 | 5.3 | 1.6 | 39.9 |
| 19 | bioclim_CHELSA_bio10_02.mean | 63.8 | 4.5 | 1.5 | 41.4 |
| 20 | terraclim_tmax_summer | 60.8 | 4.3 | 1.4 | 42.8 |
| 21 | soiltemp_fall_5_15cm | 60.7 | 3.9 | 1.4 | 44.3 |
| 22 | timetocities_mean | 57.0 | 2.9 | 1.4 | 45.6 |
| 23 | terraclim_vap_fall | 56.3 | 3.9 | 1.3 | 47.0 |
| 24 | bioclim_CHELSA_bio10_14.mean | 55.8 | 4.6 | 1.3 | 48.3 |
| 25 | soiltemp_spring_5_15cm | 55.6 | 3.5 | 1.3 | 49.6 |
| 26 | roads_sec_m_km2 | 55.6 | 3.5 | 1.3 | 51.0 |
| 27 | terraclim_tmin_fall | 51.6 | 3.7 | 1.2 | 52.2 |
| 28 | terraclim_ppt_fall | 50.4 | 4.2 | 1.2 | 53.4 |
| 29 | terraclim_ppt_spring | 49.9 | 3.7 | 1.2 | 54.6 |
| 30 | soiltemp_summer_5_15cm | 49.8 | 3.3 | 1.2 | 55.7 |
| 31 | terraclim_tmin_winter | 48.3 | 2.6 | 1.1 | 56.9 |
| 32 | terraclim_q_fall | 47.2 | 3.5 | 1.1 | 58.0 |
| 33 | terraclim_tmax_winter | 46.9 | 3.0 | 1.1 | 59.1 |
| 34 | terraclim_vap_spring | 46.6 | 2.9 | 1.1 | 60.2 |

| | | | | | |
|---|---|---|---|---|---|
| 35 | terraclim_tmax_spring | 46.6 | 2.8 | 1.1 | 61.4 |
| 36 | terraclim_aet_winter | 45.1 | 3.2 | 1.1 | 62.4 |
| 37 | bioclim_CHELSA_bio10_06.mean | 44.5 | 2.6 | 1.1 | 63.5 |
| 38 | bioclim_CHELSA_bio10_12.mean | 44.2 | 3.2 | 1.1 | 64.5 |
| 39 | roads_prim_m_km2 | 41.7 | 2.5 | 1.0 | 65.5 |
| 40 | terraclim_tmin_spring | 41.4 | 2.6 | 1.0 | 66.5 |
| 41 | terraclim_vap_winter | 41.2 | 2.3 | 1.0 | 67.5 |
| 42 | bioclim_CHELSA_bio10_07.mean | 40.2 | 1.8 | 1.0 | 68.4 |
| 43 | bioclim_CHELSA_bio10_19.mean | 40.0 | 2.2 | 1.0 | 69.4 |
| 44 | airports_distmin_m | 37.1 | 2.9 | 0.9 | 70.3 |
| 45 | bioclim_CHELSA_bio10_13.mean | 36.7 | 2.2 | 0.9 | 71.2 |
| 46 | terraclim_aet_spring | 36.7 | 2.4 | 0.9 | 72.0 |
| 47 | bioclim_CHELSA_bio10_05.mean | 35.2 | 2.1 | 0.8 | 72.9 |
| 48 | terraclim_q_winter | 34.9 | 1.8 | 0.8 | 73.7 |
| 49 | bioclim_CHELSA_bio10_09.mean | 34.4 | 1.8 | 0.8 | 74.5 |
| 50 | date | 34.0 | 2.0 | 0.8 | 75.3 |
| 51 | terraclim_soilm_fall | 33.9 | 1.4 | 0.8 | 76.1 |
| 52 | terraclim_ppt_winter | 33.6 | 1.7 | 0.8 | 76.9 |
| 53 | soiltemp_winter_5_15cm | 33.3 | 1.9 | 0.8 | 77.7 |
| 54 | lc_190 | 32.5 | 2.7 | 0.8 | 78.5 |
| 55 | terraclim_soilm_winter | 31.6 | 1.4 | 0.8 | 79.2 |
| 56 | terraclim_soilm_spring | 31.5 | 1.5 | 0.7 | 80.0 |
| 57 | bioclim_CHELSA_bio10_08.mean | 31.2 | 1.6 | 0.7 | 80.7 |
| 58 | terraclim_srad_spring | 30.6 | 1.4 | 0.7 | 81.5 |
| 59 | terraclim_soilm_summer | 30.0 | 1.3 | 0.7 | 82.2 |
| 60 | lc_shdi | 29.5 | 1.7 | 0.7 | 82.9 |
| 61 | elevation_mean | 29.4 | 1.2 | 0.7 | 83.6 |
| 62 | soilchem_ph.h2o.mean | 29.2 | 1.3 | 0.7 | 84.3 |
| 63 | soilchem_sand.wfraction.mean | 29.1 | 1.5 | 0.7 | 85.0 |
| 64 | terraclim_ws_winter | 29.0 | 1.4 | 0.7 | 85.6 |
| 65 | terraclim_ws_fall | 28.6 | 1.2 | 0.7 | 86.3 |
| 66 | terraclim_ws_summer | 28.4 | 1.4 | 0.7 | 87.0 |
| 67 | terraclim_q_spring | 28.1 | 1.1 | 0.7 | 87.7 |
| 68 | soilchem_sand.wfraction.sd | 27.4 | 1.7 | 0.7 | 88.3 |
| 69 | soilchem_ph.h2o.sd | 27.2 | 1.4 | 0.6 | 89.0 |
| 70 | soilchem_clay.wfraction.sd | 27.1 | 1.8 | 0.6 | 89.6 |
| 71 | soilchem_organic.carbon.mean | 26.8 | 1.1 | 0.6 | 90.2 |
